# ARF4-mediated Retrograde Trafficking Drives Chemoresistance in Glioblastoma

**DOI:** 10.1101/2021.07.18.451328

**Authors:** Shreya Budhiraja, Shivani Baisiwala, Ella Perrault, Sia Cho, Khizar Nandoliya, Gabriel Dara, Andrew Zolp, Li Chen, Crismita Dmello, Cheol H. Park, Adam M Sonabend, Atique U Ahmed

**Author notes:** Correspondence; The Department of Neurological Surgery, Feinberg School of Medicine, Northwestern University, 303 East Superior Street Chicago, IL60611.

## Abstract

Glioblastoma (GBM) is the most common type of adult malignant brain tumor, with a median survival of only 21 months. This is partly due to the high rate of resistance to conventional therapy, including temozolomide (TMZ), leading to recurrence rates close to 100%. It still remains unknown what drives the development of this resistance. To identify the unknown genes driving the development of this resistance, we performed a genome-wide CRISPR knockout screen comparing a DMSO-treated population with a TMZ-treated population over 14 days. We identified 4 previously unstudied genes – *ARF4*, *PLAA, SPTLC1*, and *PIGK* – that showed significant elevations in expression in recurrent tumors in patient datasets, along with significant survival benefits corresponding to low gene expression. Further investigation of *ARF4*, known to be involved in retrograde trafficking, allowed us to identify a mechanism of resistance that is mediated by increased retrograde transport of EGFR into the nucleus. Ultimately, our CRISPR-Cas9 screen has identified a promising therapeutic target, *ARF4*, which may drive GBM’s high resistance to chemotherapy.

## INTRODUCTION

Glioblastoma (GBM) is the most aggressive and common type of adult malignant brain tumor, with 12,000 new diagnoses each year [1-3]. Even with the current standard of care— surgical resection, radiation, and temozolomide (TMZ)-based chemotherapy—the median survival is still only about 20 months [4-6]. This is thought to be due to the high rate of resistance to conventional therapy, including TMZ, leading to recurrence rates close to 100% [3].

It remains largely unknown what drives the development of this fatal resistance. In an effort to determine what limits the long-term efficacy of chemotherapies like TMZ, numerous studies have compared primary and recurrent tumors. However, these results are only able to represent differences in genetic landscapes of these tumors, rather than the genes actually driving the development of this chemotherapeutic resistance. Thus, it is imperative to take a more comprehensive approach, where looking at functional effects of specific genes could uncover the major players causing this resistance.

Due to its simplicity and efficacy, CRISPR-Cas9 functional screens have been adopted in various contexts to revolutionize drug discovery and therapy [7, 8]. CRISPR (clustered regularly interspaced short palindromic repeats)-Cas9 gene modification technology utilizes (1) an RNA-guided endonuclease Cas9 that creates DNA breaks and (2) a single guide RNA (sgRNA) guides Cas9 to the appropriate gene. After the cell repairs these DNA breaks through non-homologous end-joining (NHEJ), the gene is knocked out and rendered ineffective [7].

Since the main benefit of this ambitious approach is the ability to determine which genes directly contribute to a certain phenotype before and after a specific gene’s knockout, our lab harnessed the power of this technology to elucidate mechanisms of therapeutic resistance in GBM [9]. To do so, we performed a genome-wide CRISPR knockout screen in H4 human GBM cells, encompassing over 17,000 genes. Analysis showed that there was significant enrichment in guides for known TMZ-sensitivity genes that have been highly cited—*ATG14, MSH6, MLH1*, and *PMS2*—thus validating our screen results. However, more importantly, we were able to identify a list of 200 novel genes implicated in TMZ resistance.

Of these genes, 4 previously unstudied genes—*ARF4*, *PLAA, SPTLC1*, and *PIGK*—were tested and confirmed for their protective action to TMZ, revealing potential novel mechanisms of GBM resistance to this chemotherapeutic agent. Further investigation of one particularly enriched target, *ARF4*, known for its role in regulating retrograde transport from endosomes to the trans-Golgi network (TGN), was of particular interest, since the role of retrograde trafficking is quite understudied in GBM, let alone therapeutic resistance [10-13]. In this study, we demonstrate *ARF4*’s critical role in trafficking receptors that are known to maintain GBM’s adaptive response to TMZ. In particular, we find that EGFR is more highly trafficked to the nucleus, promoting the transcription of DNA repair proteins that cause this resistance. Our study, therefore, provides evidence of a mechanism by which blocking retrograde transport reflects increased therapeutic efficacy of chemotherapeutic agents like TMZ.

## Materials & Methods

### CRISPR-Cas9 Knockout Screening

Screening was performed as previously described [14]. H4 human GBM cells were infected with the whole-genome knockout Brunello library (Addgene, Cambridge, MA, USA), which included ∼19,000 genes with 4 sgRNAs per gene and 10,000 sgRNA non-targeting controls. To prepare the library, 80% confluent HEK293T cells were harvested and seeded into a T225 flask for 20-24 h. Opti-MEM I reduced serum, psPAX −10.4 µg/ml, pMD2.G −5.2 µg/ml, and Lipofectamine plus reagent were then added to the cells. Following 4 h of incubation, the media was filtered with 0.45 µM filters. The virus was then aliquoted and stored at −80 °C.

To titer the virus, 3 million H4 cells and 2 ml of media were then seeded into a 12-well plate. After 400 ul, 200 ul, 100 ul, 75 ul, 50 ul, 25 ul of virus and 8 µg/µl of polybrene were added to each well, the cells were spinfected at 1000 g for 2 h at 33 °C. Following incubation of the cells for 24 h at 37 °C, the cells were harvested and seeded at 4000 cells per well for 96 h along with a well of non-transduced cells. Finally, a titer glo assay was conducted to determine cell viability and multiplicity of infection (MOI).

70,000 sgRNAs were used to culture and spinfect 500 million H4 cells. 150 million cells remained following 4 days of cell selection by addition of 0.6 µg/ml puromycin. 50 million of these cells were used to extract genomic DNA (gDNA) after amplification of sgRNA with unique barcoded primers that served as the control. The rest of the cells were expanded for 4 days to 200 million cells preceding treatment with 700 uM DMSO and TMZ for 14 days. The cells were then harvested in order to amplify the sgRNA and create the sequencing library.

gDNA was extracted using the Zymo Research Quick-DNA Midiprep Plus Kit (Cat no: D4075, Irvine, CA, USA) in order to amplify the sgRNA. gDNA was then cleaned with 100% ethanol with 1/10 volume 3 M sodium acetate, PH 5.2, and 1:40 glycogen co-precipitant (Invitrogen Cat no: AM9515). To measure gDNA concentration, Nano drop 2000 (Thermo Scientific, Waltham, MA, USA) was used, and PCR was performed to expand the DNA.

Next generation sequencer (Next Seq) was used to sequence the sgRNAs at 300 million reads for the four sgRNAs pool at 1000 reads/sgRNA. 80 cycles of read 1 (forward) and 8 cycles of index 1 were used to sequence the samples, as stated in the Illumina protocol. 20% PhiX was added on the Next Seq to enhance coverage.

CRISPRAnalyzR and the CaRpools pipeline were used to analyze computational data. DESeq2 and the MaGeCK algorithms were applied to determine significance of changes [15].

### Cell Lines & Culture

U251, a human glioma cell line, was obtained from the American Type Culture Collection (Manassas, VA, USA). To culture the cells, Dulbecco’s Modified Eagle’s Medium (DMEM; HyClone, Thermo Fisher Scientific, San Jose, CA, USA), was used with 10% fetal bovine serum (FBS; Atlanta Biologicals, Lawrenceville, GA, USA) and 1% penicillin-streptomycin antibiotic mixture (Cellgro, Herndon, VA, USA; Mediatech, Herndon, VA, USA).The patient-derived xenograft (PDX) glioma cells (GBM43 and GBM6), which were obtained from Dr. C. David James at Northwestern University, were cultured in DMEM with 1% FBS and 1% penicillin-streptomycin. Cells that were used for a maximum of 4 passages were replenished using a frozen stock. Frozen cells were maintained in liquid nitrogen at -180°C in pure FBS with 10% dimethyl sulfoxide (DMSO) added.

### Cellular Transfection

In order to generate lentiviral particles, low passage 293T cells (ATCC, Manassas, VA, USA) were plated at 90% confluency. After 6 hours, the cells were then transfected with a mix of HP DNA Transfection Reagent (Sigma Aldrich, St Louis, MO, USA) diluted in OptiMEM media (Gibco, Waltham, MA, USA) along with packaging and target plasmids, as noted in the manufacturer’s instructions. shRNA plasmids were procured from Genecopoeia (Rockville, MD, USA), and overexpression plasmids were obtained from AddGene (Watertown, MA, USA). After maintaining the transfected 293T cells in culture for 48-72 hours, the virus-containing supernatant was harvested, centrifuged at 1200 rpm for 5 minutes, and sterilized using a 45-micron filter.

### Viral Transduction

Following resuspension in ∼50 ul of media, ∼10-20 MOI lentivirus amounts were added per sample before adding 4ug/mL of polybrene to the virus-cell mixture. Subsequently, the virus-media-polybrene mixture was spun at 850g for 2 hours at 37 °C. After incubation overnight at room temperature, the cells were plated and maintained in culture with regular media changes for 48-72 hours. To assess the efficiency of the resulting modifications, Western blots were used.

### Animals & In Vivo Models

In this study, athymic nude mice (nu/nu; Charles River, Skokie, IL, USA) were used and were housed in compliance with Institutional Animal Care and Use Committee (IACUC) requirements along with federal and state statutes. The animals were kept in shoebox cages with food and water available along with a 12-hour light and dark cycle.

Intracranial implantation of glioblastoma cells was performed in accordance with our lab’s previously established glioblastoma mouse model. Animals first received buprenex and metacam by intraperitoneal (IP) injection. Then, they were anesthetized from a second injection of ketamine/xylazine mixture (Henry Schein; New York, NY, USA). To confirm complete sedation, mice were pinched in the foot. Artificial tears were applied to each eye for protection, and ethanol and betadine were applied to the scalp for sterilization. To expose the skull, a small incision was made using a scalpel, followed by a ∼1mm burr hole drilled above the right frontal lobe. The mice were then placed in a stereotactic rig, where a Hamilton syringe loaded with 5 uL of cell solution was used to make an injection 3 mm from the dura over a period of one minute. The needle was then raised slightly and left for an additional minute in order to ensure that the cell suspension was released. After the syringe was slowly removed, the scalp was closed with sutures while maintaining the head position (Ethicon; Cincinnati, OH, USA). The animals were postoperatively placed on heat pads until awake and reactive.

Any necessary drug treatments were started one week after the implantation, in which animals received IP injections of either TMZ (2.5 mg/kg) or equimolar DMSO. Each experimental group consisted of 5 mice with an even mix of gender. Signs of tumor progression, such as weight reduction, reduced body temperature, and hunching, were monitored for and recorded throughout the study. Animals were euthanized according to Northwestern University and IACUC guidelines following determination that they were unlikely to survive the next morning.

### Immunofluorescence

Plates were first removed from the 37 °C incubator and washed once with PBS. 200ul of 4% PFA were then added to each section for 10 minutes before cells were gently washed with PBS and blocked for 2 hours in 200ul of 10% BSA solution at room temperature. Following aspiration of BSA off of the slides, 100ul of primary antibody was mixed with 1% BSA. After cells were incubated in the 4 °C fridge overnight, they were washed 3 times for 5 minutes each in 1% BSA the next morning. 200ul of secondary antibody was then added to each section, and the plate was incubated for 2-3 hours at room temperature. Sections were washed 3 times for 5 minutes each in PBS and prolong gold anti-fade reagent with DAPi was applied to the slide. The slides were imaged using a microscope, and images were compiled and analyzed in ImageJ.

### Cell Viability Assays

Viability was assessed using the MTT assay, in which cells were plated at a density of 3000-5000 per well in a 96-well plate with 6-8 replicates for each condition. After 3x treatment with varying doses of TMZ, the media was removed and cells were treated with MTT solution. This MTT solution was made by diluting MTT stock reagent at 5mg/ml in dPBS then diluting in fresh media at a stock:media ratio of 1:10 before 110ul was added to each well. After incubation at 37 °C for 3-5 hours, the media was carefully removed without aspirating or pipetting down to prevent disturbing any crystals that had formed. Following addition of 100ul of DMSO to each well, cells were resuspended in the DMSO until the crystals dissolved, which was indicated by a color change to purple. The plate was then left at room temperature for 10 minutes before it was read on the plate reader at an absorbance of 570nm. Data analysis was performed to find percent viability in each well.

### Western Blotting

After cells were treated, detached using trypsin, and washed with PBS, they were resuspended in mammalian protein extraction reagent (M-PER; Thermo Scientific, Rockford, IL, USA) supplemented with protease and phosphatase inhibitor (PPI; Thermo Scientific, Rockford, IL, USA) and EDTA (Thermo Scientific, Rockford, IL, USA). Cells were then sonicated in a water bath 3 times for 30 seconds each with 30 second rest intervals before being centrifuged at 21,000 × g for 10 min at 4 °C. The supernatant was collected, and the protein concentration for each western blot sample was determined using BSA assays (Thermo Scientific, Rockford, IL, USA). Equal amounts of protein and varying amounts of sodium dodecyl sulfate buffer (SDS sample buffer; Alfa Aesar, Wood Hill, MA, USA) supplemented with beta-mercapto-ethanol and water were used in each sample, adding up to the same total volume. After mixing the samples, they were boiled at 95 °C for 10 minutes.

Proteins were transferred onto polyvinylidene difluoride (PVDF) membranes (Millipore, Darmstadt, Germany) using a BioRad transfer machine after being run through 10% SDS-polyacrylamide (SDS-PAGE; made in house) by gel electrophoresis using BioRad equipment (Hercules, CA, USA). The membranes were then washed 3 times in PBS for 10 mins each prior to being blocked with Tris-buffered saline (TBS) containing 0.05% Tween20 (Sigma Aldrich, St. Louis, MO, USA) and 5% powdered milk. After the membranes were blocked for 1-2 hours, they were cut for proteins of interest and placed into primary antibody solutions consisting of the appropriate ratio of antibody to 5% BSA solution supplemented with sodium azide. Membranes were incubated overnight on a shaker at 4°C before being washed and incubated in secondary antibody diluted 1:4000 in 5% milk. Finally, membranes were washed 3 times for 20 minutes each, coated with enhanced chemiluminescence (ECL; Clarity ECL, BioRad), and developed using a developer machine (BioRad, Hercules, CA, USA).

### Flow Cytometry

To measure ER stress, cells were transfected with the appropriate ER stress reporter (Addgene, Cambridge, MA, USA). After 48-72 hours, cells were washed once with PBS, harvested, resuspended in 80 μl of PBS, and transported to the flow cytometry core in darkness. Cells were gated for live cells, single cells, and ER reporter positivity on the Fortessa Analyzer. Analysis was performed in FlowJo.

### Single Cell RNA-Seq

Single cell Drop-Seq was performed according to previously published protocols [16]. After cells were isolated, a single cell suspension was formed and the sample was run through a Drop-Seq set up before sequencing. Analysis was performed using the Seurat pipeline [17].

Single cell data on gene localization in GBM was acquired from the publically available data browser at gbmseq.org [18].

### Quantitative PCR

RNA was first extracted using Qiagen RNEasy kits (Qiagen, Hilden, Germany). cDNA was then created from the RNA samples using an iScript kit (BioRad, Hercules, CA, USA). After diluting cDNA 1:10 in distilled water, reactions were set up in triplicates with standard amounts of cDNA, SyberGreen (Thermo Fisher, Rockford, IL, USA), and forward and reverse primers (IDT, Newark, NJ, USA) to use for downstream quantitative PCR. Results were read out on a standard qPCR machine. All primers were generated using Primer-BLAST.

### Statistical Analysis

GraphPad Prism v8.0 software (GraphPad Software; San Diego, CA, USA) was used to conduct statistical analyses. Overall, data are presented as mean with standard deviation for continuous variables and number or percentage for categorical variables. Student’s t test or Wilcoxon rank sum test was used to assess differences between two groups, while ANOVA with post hoc Tukey’s test or Mann–Whitney U test followed by Bonferroni correction was used to assess differences among multiple groups. Survival curves were graphed with the Kaplan-Meier method and compared by log-rank test. For statistical significance, p < 0.05 was used, and all tests were two-sided. Experiments including Western blots, fluorescence activated cell sorting analysis (FACS), and microarray were conducted in biological triplicate. For in vivo experiments, each group contained at least five animals, with equal representation of males and females. Each animal was treated as a technical replicate.

## Results

### Whole-genome CRISPR Sensitivity Screen Reveals Genes Conferring Resistance to Chemotherapy in GBM

In order to identify novel genes implicated in TMZ resistance, an unbiased whole-genome CRISPR-Cas9 knockout screen was performed in human H4 GBM cells. Involving more than 19,000 genes and 2000+ non-targeting controls, this screen utilized the whole-genome knockout Brunello library, with a coverage of 4 sgRNAs per gene. For 14 days, a DMSO-treated population was compared to a TMZ-treated population, with the TMZ-treated population receiving 700 uM of TMZ every other day. After the 14 days, cells were isolated and DNA libraries were prepared, and samples from the DMSO-treated and TMZ-treated conditions were sequenced for resulting gene counts to be analyzed **(Fig 1A)**.

**Figure 1:**
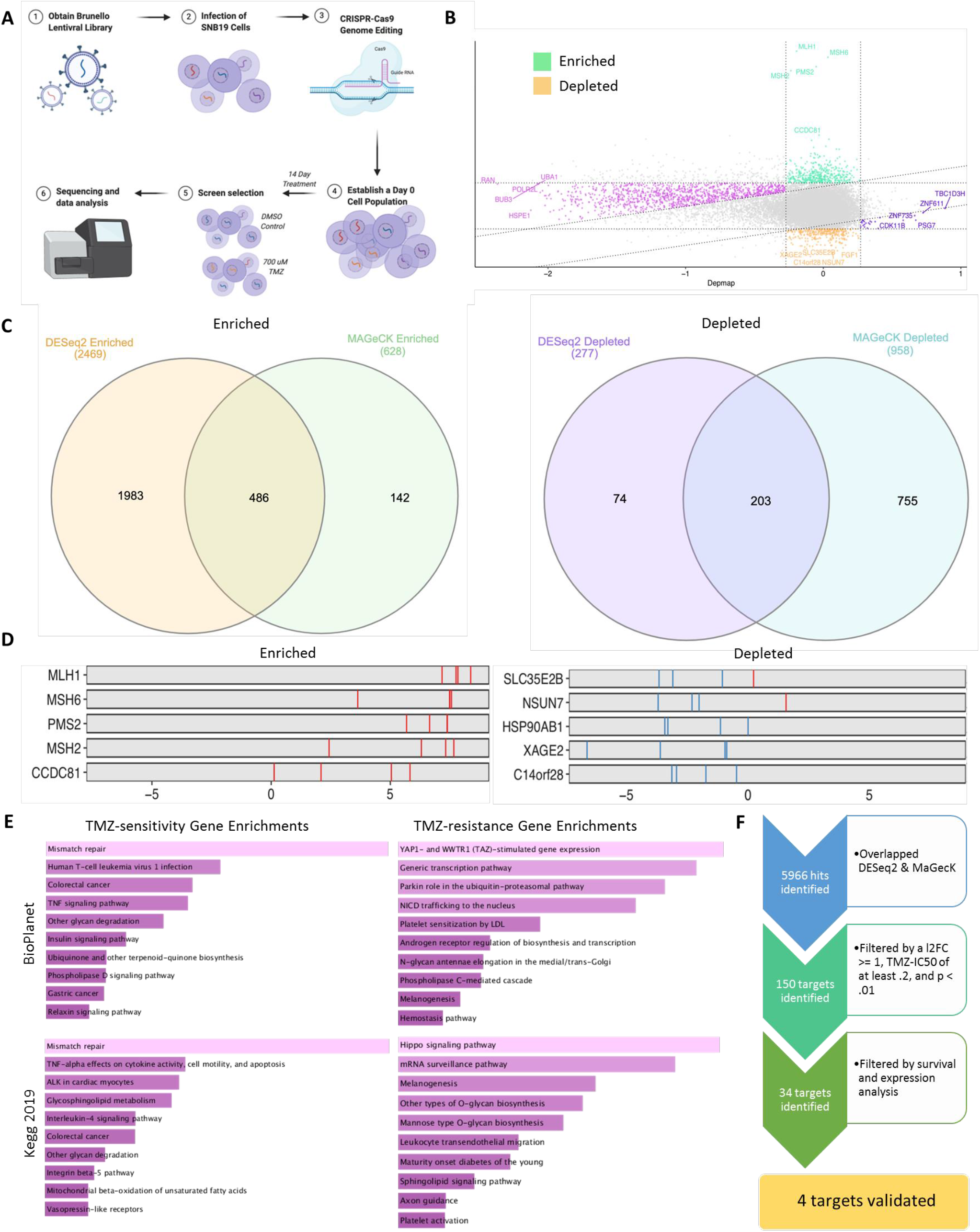
Whole-genome CRISPR Sensitivity Screen Reveals Genes Conferring Resistance to Chemotherapy in GBM. ***A)*** CRISPR-Cas9 knockout screening was performed using the Brunello whole-genome library, covering 19,000+ guides, ∼2000 non-targeting controls, and 4 sgRNAs per gene. A DMSO-treated population was compared to a TMZ-treated population, and guides were sequenced at d0 and d14. ***B)*** Comparison of d14 versus day 0 revealed segments of guides that were enriched and depleted. ***C)*** TMZ-sensitivity and TMZ-resistance genes identified and pursued were filtered with multiple CRISPR-appropriate analysis algorithms ***D)*** Specific genes were then examined for enrichment and depletion at day 0 and day 14. Known TMZ-resistance genes were depleted at d14, reflecting the validity of our screen. Key TMZ-sensitivity genes were identified as enriched (*MLH1, MSH6, PMS2, MSH2*), further reflecting validity of the screen. ***E)*** Enrichment mapping identified key pathways across identified TMZ-sensitivity and TMZ-resistance genes. Enrichment mapping was performed in the Enrichr application. ***F)*** Of the 5966 genes identified as TMZ-resistance genes, 150 were significant by DESeq2 and MAGeCK algorithms. 34 were filtered based on novelty, ability to study, and clinical significance. 4 were finally chosen for validation to reflect critical pathways from the screen. Analysis was performed in Prism 8, using ANOVA to compare row-means to determine significance or using log-rank tests to determine survival significance *p < 0.05; **p < 0.01; ***p < 0.001; ****p < 0.0001; ns, not significant.

Over the course of our screen, guides were either enriched or depleted. Guides corresponding to genes involved in conferring resistance to TMZ, or TMZ-resistance genes, were depleted, whereas those corresponding to TMZ-sensitivity genes were enriched **(Fig 1B)**. Genomic sequencing quality evaluations achieved high per-base sequence quality, an excellent quality score distribution over sequence length, and an expected length distribution for all sequences **(Fig S1A)**. Furthermore, the cumulative number of sgRNAs exhibited different distributions across DMSO and TMZ experimental conditions, with the number of sgRNAs in the TMZ condition at d14 experiencing depletion **(Fig S1B, S1C)**. This is consistent with the expectation that TMZ treatment would lead to a depletion of guides for TMZ-resistance genes, while DMSO treatment would represent a control condition with most sgRNA’s present.

After applying different analysis methods, including DESeq2 and MAGeCK, between the 2 conditions, we were able to identify several targets for our study. Any gene that appeared in at least two of the analysis methods indicated a gene of interest **(Fig 1C)**. From this selection criteria, we were able to employ a critical positive control for known TMZ-sensitivity and TMZ-resistance genes. We found that known TMZ-resistance genes were significantly depleted, validating the reliability of our screen results. We then found that the TMZ-sensitivity genes involved in mismatch repair –– including *MLH1, MSH6, PMS2, MSH2* –– were highly enriched, further validating the accuracy of our screen **(Fig 1D)** [19-21]. We were also able to perform enrichment analysis on the most enriched pathways in both conditions. The resulting pathways of interest involved *NICD* trafficking and Hippo signaling pathways for TMZ-resistance, along with mismatch repair and *IL-4* signaling pathways for TMZ-sensitivity **(Fig 1E)**.

Again, the goal of our study was to identify which genes are involved in driving therapeutic resistance to TMZ, as these genes represent novel therapeutic targets. To identify these potential targets, we selected TMZ-resistance genes – corresponding to depleted guides – that demonstrated significant log fold change in our screen, were linked to significant survival differences in patient datasets, and mirrored pathways that were globally enriched across our sample. Additionally, the target genes were chosen for their novelty within the study of GBM and showed diversity among the pathways they were implicated in. Initially, 150 TMZ-resistance genes were identified, but ultimately 4 genes of interest were chosen: *ARF4, PLAA, SPTLC1*, and *PIGK* **(Fig 1F)**.

### Identified Genes Show Effects on Patient Survival

We first examined publicly available GBM patient datasets to understand how expression of our genes of interest varied across different conditions and how this expression was linked to survival data. This initially helped identify our 4 target genes from our larger set of TMZ-resistance genes. Analysis of CGGA (Chinese Glioma Genome Atlas) data through the GlioVis portal showed that *ARF4* and *SPTLC1* mRNA expression was especially elevated in GBM compared to non-tumor *(p<0.001)*, while *PIGK* and *PLAA* mRNA expressions were similar between the two conditions **(Fig 2A)**.

**Figure 2:**
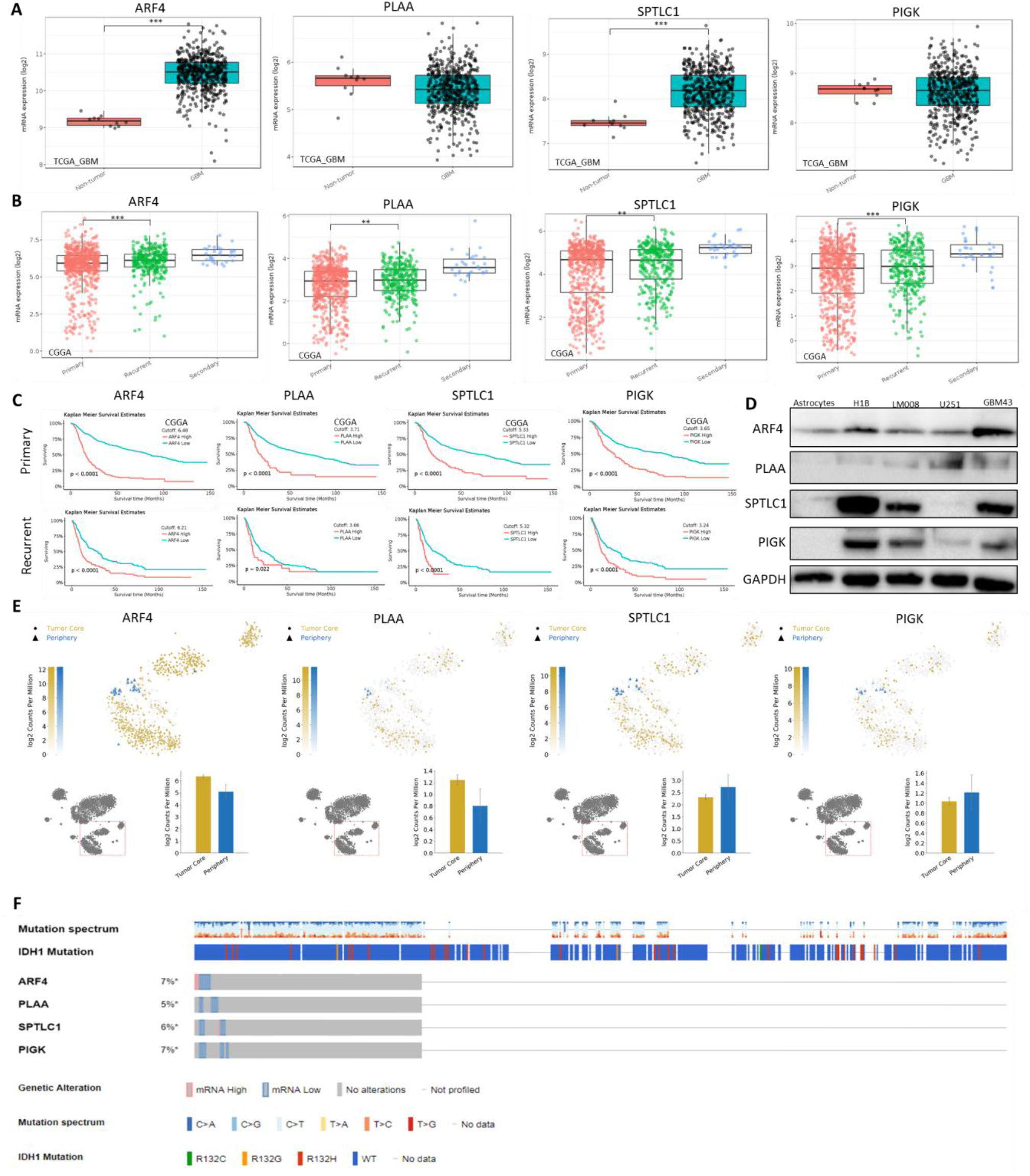
Bioinformatic analysis reveals that four identified genes have effects on patient survival. ***A)** ARF4, PLAA, SPTLC1*, and *PIGK* mRNA expression levels were analyzed using the Cancer Genome Atlas (TCGA) for non-tumor and GBM patients. GBM patients had higher mRNA expression levels of all four genes than non-tumor patients. ***B)*** *ARF4, PLAA, SPTLC1*, and *PIGK* mRNA expression levels were analyzed using the Chinese Cancer Genome Atlas (CCGA) for patients with primary, secondary, and recurrent tumors. Patients with recurrent tumors had higher mRNA expression levels of all four genes than patients with primary tumors. ***C)*** All GBM patients were stratified into up-regulated and down-regulated groups based on gene expression using quartile (Q1, Q3) as split points. High expression of *ARF4, PLAA SPTLC1*, and *PIGK* correlated with reduced median survival in patients with primary and recurrent tumors. Survival curves were generated via the Kaplan-Meier method and compared by log-rank test, using optimal cutoff analyses. ***D)*** Immunoblot analysis of *ARF4, PLAA, SPTLC1*, and *PIGK* expression in astrocytes, H1B.F3 (NSC), LM008 (NSC), U251, and PDX line GBM43. Protein extract of these lines were immunoblotted with antibody against *ARF4, PLAA, SPTLC1*, and *PIGK* or an antibody against GAPDH as a control for equal loading. ***E)*** GBMSeq (publically available single-cell RNA-sequencing database) was employed to determine the location of *ARF4, PLAA, SPTLC1*, and *PIGK* in glioblastoma samples. A tSNE representation of the neoplastic cells colored based their location in the tumor core or periphery along with the gradient of color reveals where each gene is expressed. ***F)*** cBioPortal data reveals the mutation profiles of *ARF4, PLAA, SPTLC1*, and *PIGK*, where each column represents an individual patient. Analysis was performed in Prism 8, using ANOVA to compare row-means to determine significance or using log-rank tests to determine survival significance *p < 0.05; **p < 0.01; ***p < 0.001; ****p < 0.0001; ns, not significant.

We then wanted to probe the patient dataset in order to investigate the relationship between the mRNA expression of our genes of interest and recurrence status. Since our targets were classified as TMZ-resistance genes from our screen, we expected that our genes of interest would display higher mRNA expression in recurrent tumors than in primary tumors. Results showed higher mRNA expression for all genes of interest in recurrent tumors when compared to primary tumors *(p<0.01)* **(Fig 2B)**. Next, to ascertain the clinical relevance of these targets, we correlated patient survival to mRNA expression of our genes of interest. Analysis of the patient data revealed that high expression of all four targets was associated with significantly poorer survival of primary GBM patients *(p<0.001)*. More importantly, these genes echoed this trend more robustly in patients with recurrent GBM, highlighting the significance of these genes in promoting therapeutic resistance *(p<0.05)* **(Fig 2C)**.

Our next step was to examine protein expression of these four genes through western blotting in order to better understand the baseline expression levels in multiple cell lines. Western blot analysis revealed that these genes were generally expressed more in U251 and patient-derived xenograft (PDX) GBM line GBM43 than in astrocytes or normal neural stem cells (NSCs) H1B.F3 and LM008 **(Fig 2D)**. Furthermore, this expression was varied among GBM cell lines, as also seen at the RNA level, indicating that each gene’s role in developing resistance may be dependent on subtype-specific processes **(Fig S2A)**. Furthermore, mRNA expression of all of our genes of interest was found to vary among glioma types **(Fig S2B, S2C)**. These results led us to perform all validation experiments in multiple cell lines going forward.

We finally wanted to examine the location of gene expression in the tumor as well as mutation profiles. Publicly available RNA-Sequencing data gathered from GBMSeq indicated that both *ARF4* and *PLAA* were expressed more highly in the tumor core, while *SPTLC1* and *PIGK* were expressed more highly in the tumor periphery **(Fig 2E)**. Additionally, mutation profiles gathered from cBioPortal showed that all genes of interest did not have high rates of mutations in GBM, with each ranging from 5-7% **(Fig 2F)**.

### Identified Genes are Elevated during TMZ Treatment

In order to validate the results of our screen, we first performed *in vitro* experiments to confirm that our genes of interest were truly elevated during TMZ therapy. Preliminary experiments waiting 24h and 48h after 50 uM of TMZ was administered to cells yielded no upregulation at the RNA or protein levels **(Fig S3A, S3B)**. Additionally, treatment of TMZ for a total or one or two exposures (2 days apart) similarly did not result in an upregulation of our genes of interest at RNA and protein levels **(Fig S3C, S3D)**. Because of these results, we developed a model of multiple exposures of treatment, in which cells were either treated with 50 uM TMZ or equimolar vehicle control DMSO every two days for a total of three exposures **(Fig 3A)**. This model was used in order to ensure that the GBM cells undergoing TMZ treatment would exhibit a TMZ-resistant phenotype consistent with therapy that patients would typically undergo. After cells were treated in this manner, we first assessed mRNA expression in two cell lines and discovered that multiple TMZ doses resulted in substantial elevations of our genes of interest when compared to DMSO conditions **(Fig 3B)**. We then verified this trend at the protein level and reported that multiple TMZ treatments similarly resulted in increased target expression relative to the control in all three cell lines **(Fig 3C)**. This thus confirmed that the effect of TMZ-resistance gene upregulation is noted when cells are forced into resistance from multiple exposures of TMZ.

**Figure 3:**
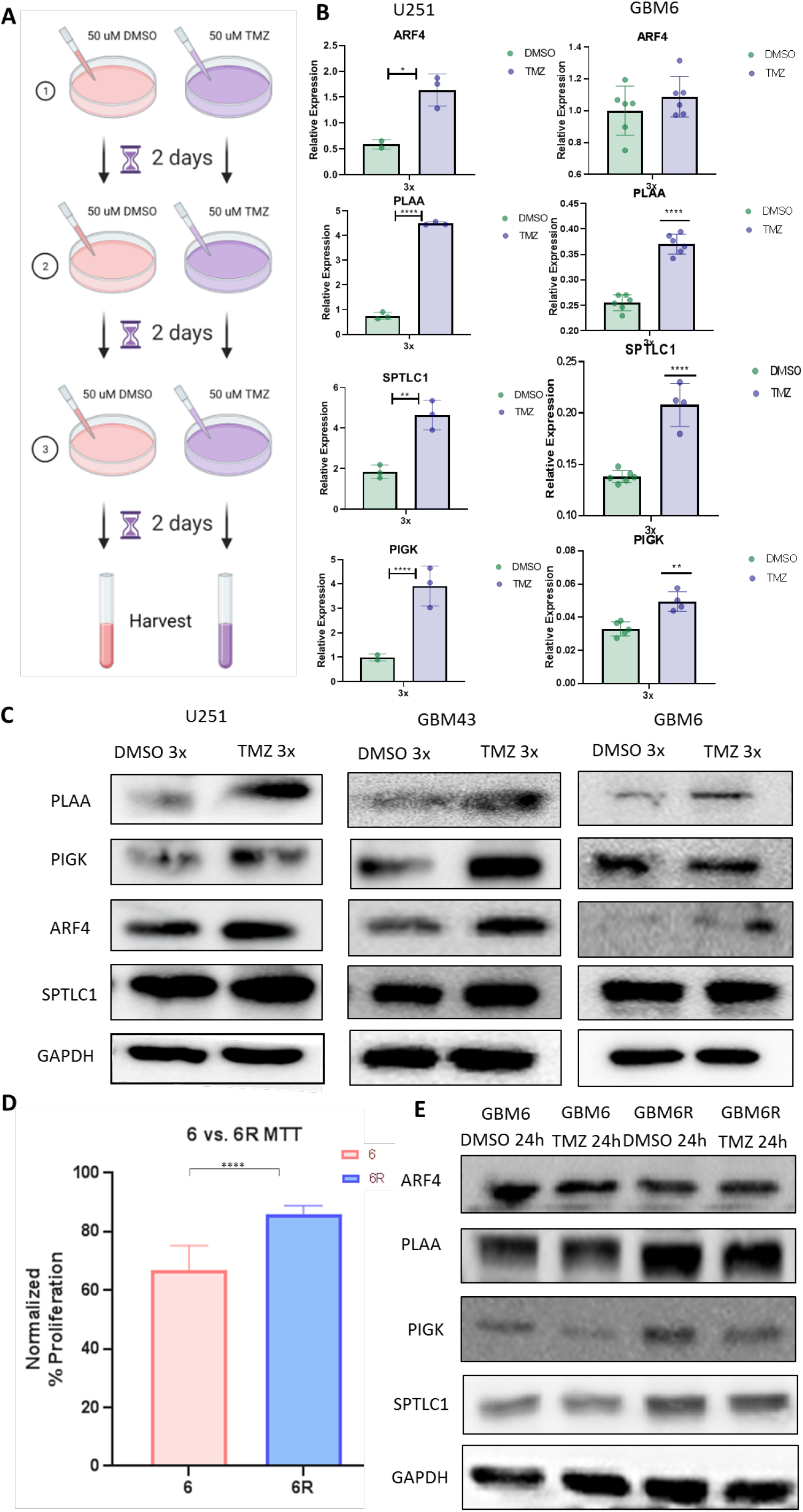
Identified Genes are Elevated during TMZ Treatment. ***A)*** A multiple-exposure model was established to generate a phenotype where cells are forced into resistance. ***B)*** *ARF4, PLAA, SPTLC1*, and *PIGK* mRNA expression levels were analyzed in 3x-treated DMSO and TMZ U251 and GBM43 cells using quantitative real-time PCR (qPCR). ***C)*** Immunoblot analysis of *ARF4, PLAA, SPTLC1*, and *PIGK* protein expression in 3x-treated DMSO and TMZ U251, GBM43, and GBM6 cells. Protein extract of these lines were immunoblotted with antibody against *ARF4, PLAA, SPTLC1, and PIGK* or an antibody against *GAPDH* as a control for equal loading. ***D)*** Proliferation GBM6 and GBM6R was assessed using MTT assays after treatment 3x treatment of 50 uM TMZ. GBM6R is confirmed to display resistance to TMZ. ***E)*** Immunoblot analysis of *ARF4, PLAA, SPTLC1*, and *PIGK* protein expression in GBM6 and GBM6R cells treated with DMSO or TMZ. Protein extract of these lines (after 24 hours of treatment) were immunoblotted with antibody against *ARF4, PLAA, SPTLC1, and PIGK* or an antibody against *GAPDH* as a control for equal loading. Analysis was performed in Prism 8, using ANOVA to compare row-means to determine significance or using log-rank tests to determine survival significance *p < 0.05; **p < 0.01; ***p < 0.001; ****p < 0.0001; ns, not significant.

Finally, we were able to determine the expression of these genes in a therapy-resistant glioblastoma (GBM6R) from a PDX of primary glioblastoma (GBM6) that underwent TMZ treatment in order to develop stable resistance to TMZ. GBM6R cells displayed a significant decrease in sensitivity to TMZ in vitro when treated multiple times with 50 uM TMZ *(p<0.0001)* and thus, we believed it to be an appropriate cell line to determine the expression of our target TMZ-resistance genes **(Fig 3D)**. After 24h of TMZ treatment in both GBM6 and GBM6R cells, our targets displayed higher protein expression across all GBM6R conditions when compared to GBM6 conditions, further validating that our genes of interest are associated with a resistant phenotype **(Fig 3E)**.

### Identified Genes Show Effects on TMZ Resistance

Because our CRISPR-Cas9 screen utilized knockouts to assess the function of each gene during the TMZ treatment, it was necessary to conduct a series of *in vitro* knockdown experiments on our genes of interest to properly confirm whether they were contributing to the TMZ resistance. To investigate the effect of knockouts on GBM cell survival after several TMZ exposures, knockdown cell lines *ARF4, PLAA, SPTLC1*, and *PIGK* were generated in U251, GBM43, and GBM6 cell lines using shRNA plasmids **(Fig 4A)**. The efficiency of three shRNA plasmids per gene were evaluated by western blotting, and the shRNAs that yielded the most significant knockdowns were adopted in the remaining experiments **(Fig S3E)**.

**Figure 4:**
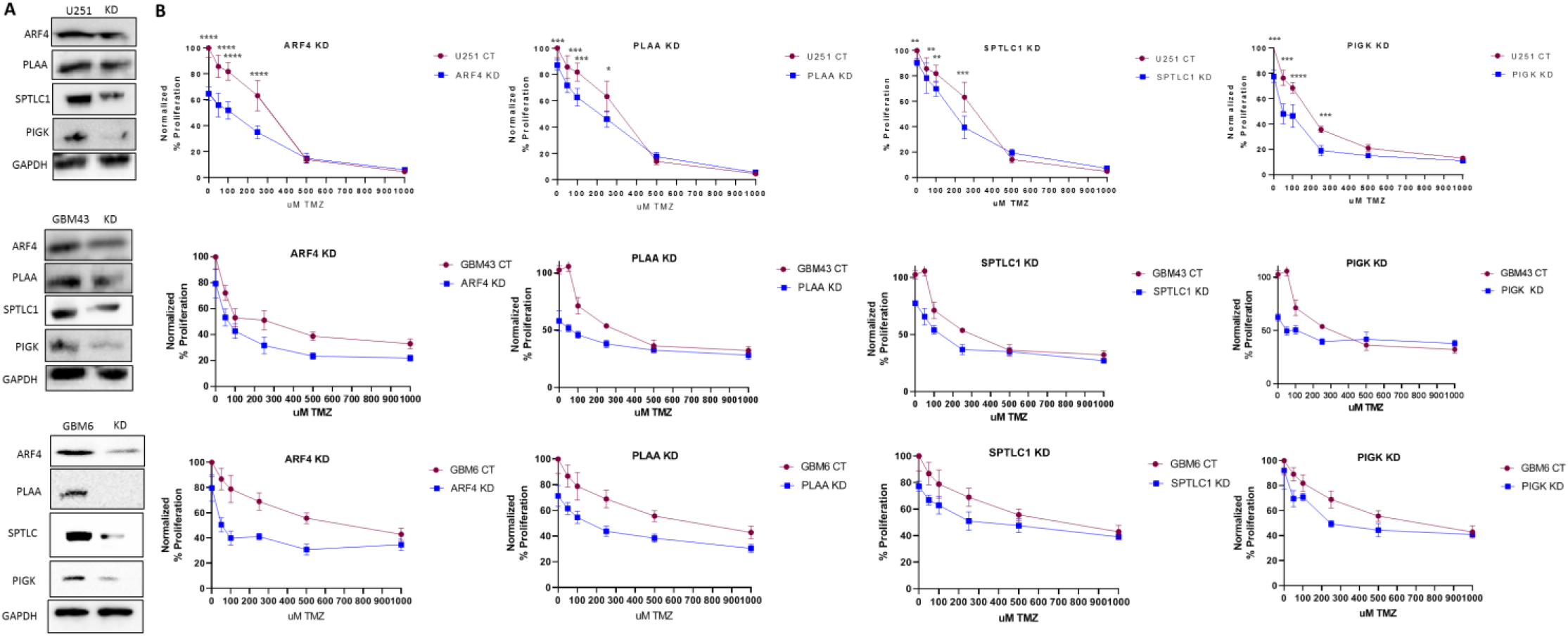
Identified Genes Show Effects on TMZ Resistance. ***A)*** Lentiviral knockdowns using CRISPR-Cas9 constructs were established in multiple cell lines – U251, GBM43, and GBM6 – and were assessed by western blotting. ***B)*** Viability of cells in multiple cell lines was assessed after three exposures of TMZ at varying doses. Cells with *ARF4, PLAA, SPTLC1*, and *PIGK* knockdowns exhibited reduced proliferation during TMZ treatment compared to control cells, showing that these genes are responsible for TMZ-resistance. Analysis was performed in Prism 8, using ANOVA to compare row-means to determine significance or using log-rank tests to determine survival significance *p < 0.05; **p < 0.01; ***p < 0.001; ****p < 0.0001; ns, not significant.

After establishing the knockdowns, an MTT assay was performed in order to determine the viability of U251, GBM43, and GBM6 cells with knockdowns of our genes of interest when supplied with multiple exposures of TMZ. Since we identified these genes as essential to conferring resistance to TMZ, we expected that knockdown of these genes would result in heightened sensitivity to TMZ and thus greater cell death. Indeed, for all knockdowns – *ARF4, PLAA, SPTLC1*, and *PIGK* – we found that GBM cells showed significant reduction in viability when compared to control cells after three treatments of TMZ **(Fig 4B)**. This thus confirmed that when our genes of interest are knocked down, they are unable to shield GBM cells with protection against TMZ.

To further validate the importance of the role of the genes of interest during therapy, single cell RNA sequencing was performed. RNA expression in mice isolated post-DMSO treatment, post-TMZ treatment, and during TMZ treatment was determined for *ARF4, PLAA, SPTLC1*, and *PIGK* **(Fig S4A)**. RNA expression for all genes was found to be especially elevated in the middle timepoint during therapy, corroborating its critical role in promoting resistance **(Fig S4B, S4C, SC)**.

### TMZ-induced ER Stress Leads to Upregulation of ARF4 and Retrograde Trafficking of Receptor Tyrosine Kinases (RTKs)

*ARF4*, or ADP-ribosylation factor 4, is a GTP-binding protein that plays a key role in the retrograde endomembrane trafficking of proteins [22-24]. When proteins or receptors need to be internalized in order to take effect in the cell, cargo is packaged into membranous structures in order to be trafficked from endosomes to the trans-Golgi network [10-13]. The co-localization of the Golgi apparatus and *ARF4* is visualized using immunocytochemistry **(Fig S5A)**. Thus, regulation of the retrograde trafficking pathway is essential for maintaining cellular homeostasis and efficient protein localization [25, 26]. As a result, defective retrograde trafficking has been linked to several malignancies, including GBM, where any alteration in protein trafficking may modulate protein levels in order to promote chemoresistance [27-30].

Given the importance of the retrograde trafficking in cancer and our identification of *ARF4* as a critical target in our screen, we postulated that TMZ is upregulating *ARF4* to accelerate the shuttling of chemoresistance promoting receptor tyrosine kinases (RTKs) into the nucleus **(Fig 5A)**. First, we sought to clarify why *ARF4* was upregulated following TMZ therapy. Previous research has demonstrated that ER stress induces *ARF4* overexpression via a number of ER and endosomal shock-related pathways [31-33]. In order to test this, we performed FACS to identify the percentage of cells with phosphorylated-*PERK*, or protein kinase RNA-like endoplasmic reticulum, that is induced via Unfolded Protein Response (UPR) pathway – a known indicator of ER stress – during TMZ treatment [34, 35]. Across two cell lines – U251 and GBM43 – the percentage of ER stress positive cells was higher when cells were given 50 uM TMZ as compared to DMSO *(p<0.001)*; more specifically, three exposures of TMZ resulted in a dramatic increase in ER stress in U251s *(p<0.0001)* **(Fig 5B)**. This supports our previous findings that cells are only forced into a state of resistance after multiple exposures of TMZ and that multiple exposures of TMZ are needed to upregulate *ARF4* significantly. In order to strengthen our evidence of this TMZ-induced ER stress response, we explored the protein expression of known ER stress markers *Calnexin, IRE1*, and *GRP78* through western blotting [35, 36]. Comparing three times DMSO-treated and TMZ-treated GBM cells in two cell lines – GBM43 and GBM6 – we were able to see elevated levels of these markers in TMZ conditions **(Fig 5C)**. Again, we were also able to see a co-upregulation of *ARF4*, suggesting that TMZ is able to increase *ARF4* expression due to its ability to trigger ER stress **(Fig 5C)**.

**Figure 5:**
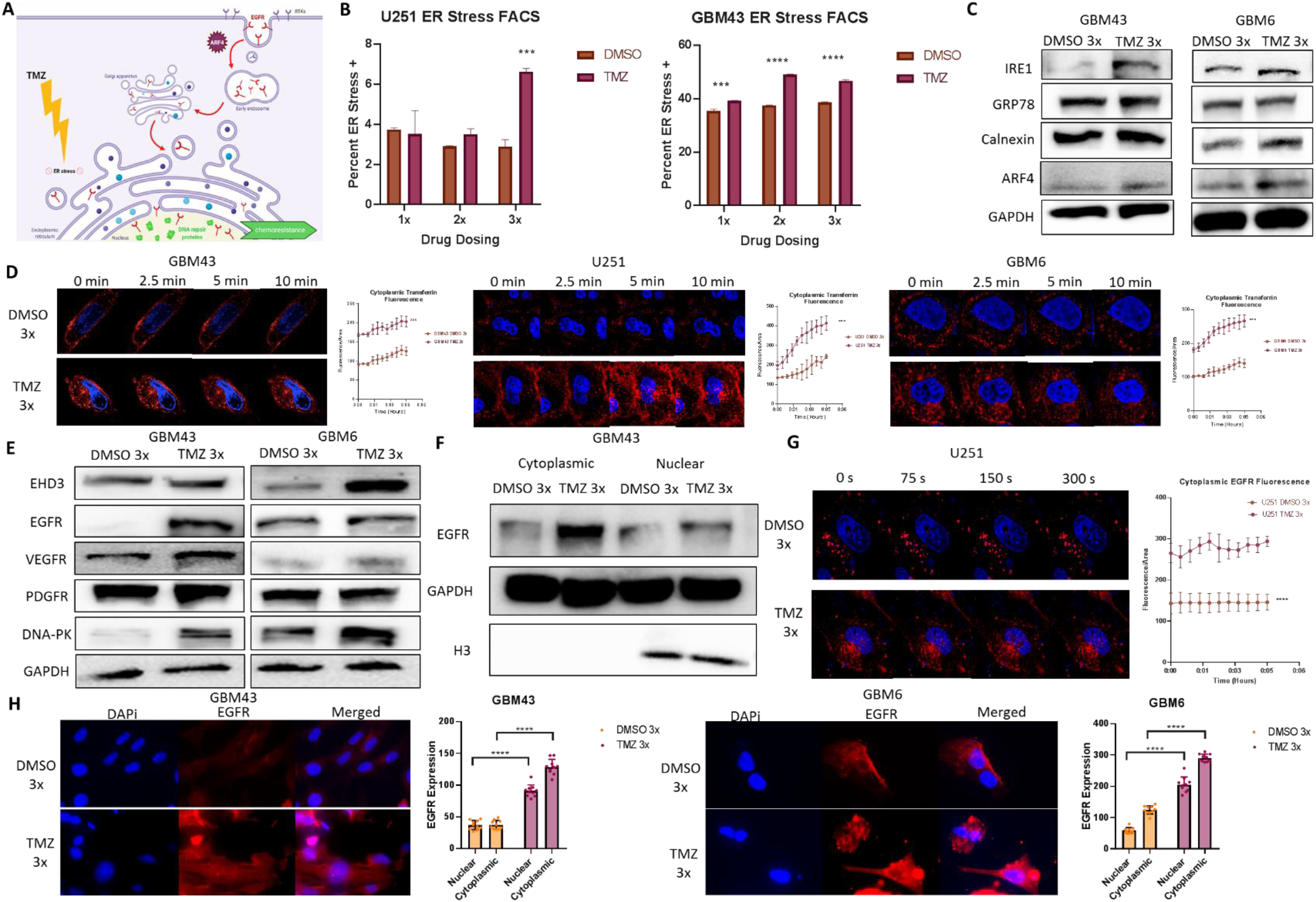
TMZ-induced ER Stress Leads to Upregulation of ARF4 and Retrograde Trafficking of Receptor Tyrosine Kinases (RTKs). ***A)*** Schematic of hypothesis. TMZ is upregulating ARF4 via ER stress, thereby increasing retrograde trafficking of resistance-promoting receptor tyrosine kinases (RTKs) like EGFR into the nucleus, transcriptional activating DNA-PK. ***B)*** FACS analyses were performed to determine how TMZ treatment alters ER stress levels in U251 and GBM43 cell lines. Samples were analyzed after 1x, 2x, or 3x treatment with 50 uM DMSO or TMZ. ***C)*** Immunoblot analysis of *IRE1* (ER stress marker)*, GRP78* (ER stress marker)*, Calnexin* (ER stress marker), and *ARF4* protein expression in 3x-treated DMSO and TMZ GBM43 and GBM6 cells. Protein extract of these lines were immunoblotted with antibody against *IRE1, GRP78, Calnexin*, and *ARF4* or an antibody against *GAPDH* as a control for equal loading. ***D)*** Live cell imaging of transferrin receptor after 0, 2.5, 5, and 10 minutes of exogenous introduction in 3x-treated DMSO and TMZ GBM43, U251, and GBM6 cells, along with quantitative analysis of cytoplasmic fluorescence over time. ***E)*** Immunoblot analysis of *EHD3* (retrograde transport marker)*, EGFR* (RTK)*, VEGFR* (RTK), and *PDFGR* (RTK), and *DNA-PK* (DNA repair protein) protein expression in 3x-treated DMSO and TMZ GBM43 and GBM6 cells. Protein extract of these lines were immunoblotted with antibody against *EHD3, EGFR, VEGFR, PDGFR*, and *DNA-PK* or an antibody against *GAPDH* as a control for equal loading. ***F)*** Immunoblot analysis of *EGFR* in cytoplasmic and nuclear fractions of 3x-treated DMSO and TMZ GBM43 cells. Protein extract of these lines were immunoblotted with antibody against *EGFR* or an antibody against *GAPDH* as a control for equal loading of cytoplasmic protein and *H3* for nuclear protein. ***G)*** Live cell imaging of EGFR after 0, 75, 150, and 300 seconds of EGF introduction in 3x-treated DMSO and TMZ GBM6 cells, along with quantitative analysis of cytoplasmic fluorescence over time. ***H)*** Immunocytochemistry of EGFR in 3x-treated DMSO and TMZ GBM43 and GBM6 cells, along with quantitative analysis of nuclear vs. cytoplasmic fluorescence. Analysis was performed in Prism 8, using ANOVA to compare row-means to determine significance or using log-rank tests to determine survival significance *p < 0.05; **p < 0.01; ***p < 0.001; ****p < 0.0001; ns, not significant.

After explaining that *ARF4*’s upregulation is due to TMZ-induced ER stress, we then wanted to confirm that *ARF4*’s role in retrograde trafficking is being altered during this treatment. To do so, we utilized live cell imaging, a powerful technique that can record intracellular activity – like retrograde trafficking – in real time [37, 38]. To identify changes in retrograde trafficking in cells treated with TMZ as compared to DMSO, we recorded a widely-used retrograde trafficking marker, the transferrin receptor (*TfR*) [39, 40]. Results from three cell lines – U251, GBM43, and GBM6 – showed that exogenous *TfR* is visibly internalized and trafficked into the cell at much higher level via the retrograde transport route in the TMZ conditions *(p<0.001)* **(Fig 5D)**.

Given the drastic alterations in global receptor trafficking seen via live cell imaging, we performed western blotting to determine if transport of known receptor tyrosine kinases (RTKs) is similarly being shuttled into GBM cells at greater rates during TMZ therapy. Western blot analysis comparing DMSO-treated and TMZ-treated cells confirmed this hypothesis, where TMZ-conditions showed higher expression of retrograde transport marker *EHD3* and known RTKs – *EGFR, VEGFR*, and *PDGFR* – in TMZ conditions **(Fig 5E)** [41]. Interestingly, we also see that *DNA-PK*, a DNA repair protein that is known to be transcriptionally activated by *EGFR*, is also more highly expressed in TMZ conditions [42, 43].

We decided to focus on *EGFR*, a known chemoresistance-promoting agent, due to multiple studies showing that *EGFR* is important for GBM survival and adaptability through its ability to activate DNA repair proteins like *DNA-PK*, especially since TMZ is known be effective against GBM cells due to its role in creating DNA damage **(Fig S5B)** [42-44]. To confirm the specific localization of *EGFR*, western blotting after nuclear and cytoplasmic fractionation displayed significant increases in *EGFR* expression in the nucleus in the TMZ condition, suggesting that this TMZ-induced alteration of the retrograde trafficking pathway may enhancing nuclear *EGFR* translocation **(Fig 5F)**. This western blotting also conveyed increased expression of *EGFR* in the cytoplasmic compartments, as explained by greater internalization of receptors similar to *TfR* and an independent mechanism of *EGFR* upregulation **(Fig 5F)** [45, 46].

To confirm this modification of *EGFR* trafficking, we then visualized endogenous *EGFR* transport via the retrograde pathway in both DMSO and TMZ conditions. Live cell imaging revealed that TMZ conditions are associated with much higher EGFR expression as well as dynamics *(p<0.0001)* **(Fig 5G)**. Immunocytochemistry of 3x DMSO- and TMZ-treated GBM43 and GBM6 cells verified this increased nuclear and cytoplasmic expression of EGFR in TMZ conditions *(p<0.0001)*, further strengthening our evidence that this TMZ-upregulated retrograde trafficking causes high nuclear translocation of *EGFR* **(Fig 5H)**.

Ultimately, we have shown that TMZ causes an upregulation of retrograde trafficking of many receptor tyrosine kinases in general. One in particular, *EGFR*, is notorious for promoting chemoresistance and is already upregulated via TMZ through an independent mechanism. Here, we see that TMZ treatment leads to dysfunctional retrograde trafficking that may leverage increased nuclear *EGFR* trafficking to strengthen the DNA repair response associated with recurrence phenotypes [42, 43].

### ARF4-mediated Retrograde Trafficking of EGFR to the Nucleus Promotes Chemoresistance through Transcriptional Activation of DNA-PK

We then wanted to confirm that this TMZ-associated upregulation of retrograde trafficking and *EGFR* nuclear translocation is an *ARF4*-dependent mechanism. To do so, we compared *TfR* trafficking in control and *ARF4*-knockdowns in multiple cell lines – U251, GBM43, and GBM6 – all of which revealed significantly reduced trafficking *(p<0.01)*, as illustrated by the lack of internalization of *TfR* that consistently stays packed to the membrane over time **(Fig 6A)**. In contrast, *ARF4*-overexpression cells resulted in significantly increased trafficking as seen by higher cytoplasmic fluorescence over time *(p<0.0001)* **(Fig 6A)**. This result also confirmed that overexpressing *ARF4* in GBM cells is able to invoke a phenotype similar to TMZ-treated GBM cells, where the increased retrograde trafficking activity is explained through ER stress-mediated *ARF4* upregulation.

**Figure 6:**
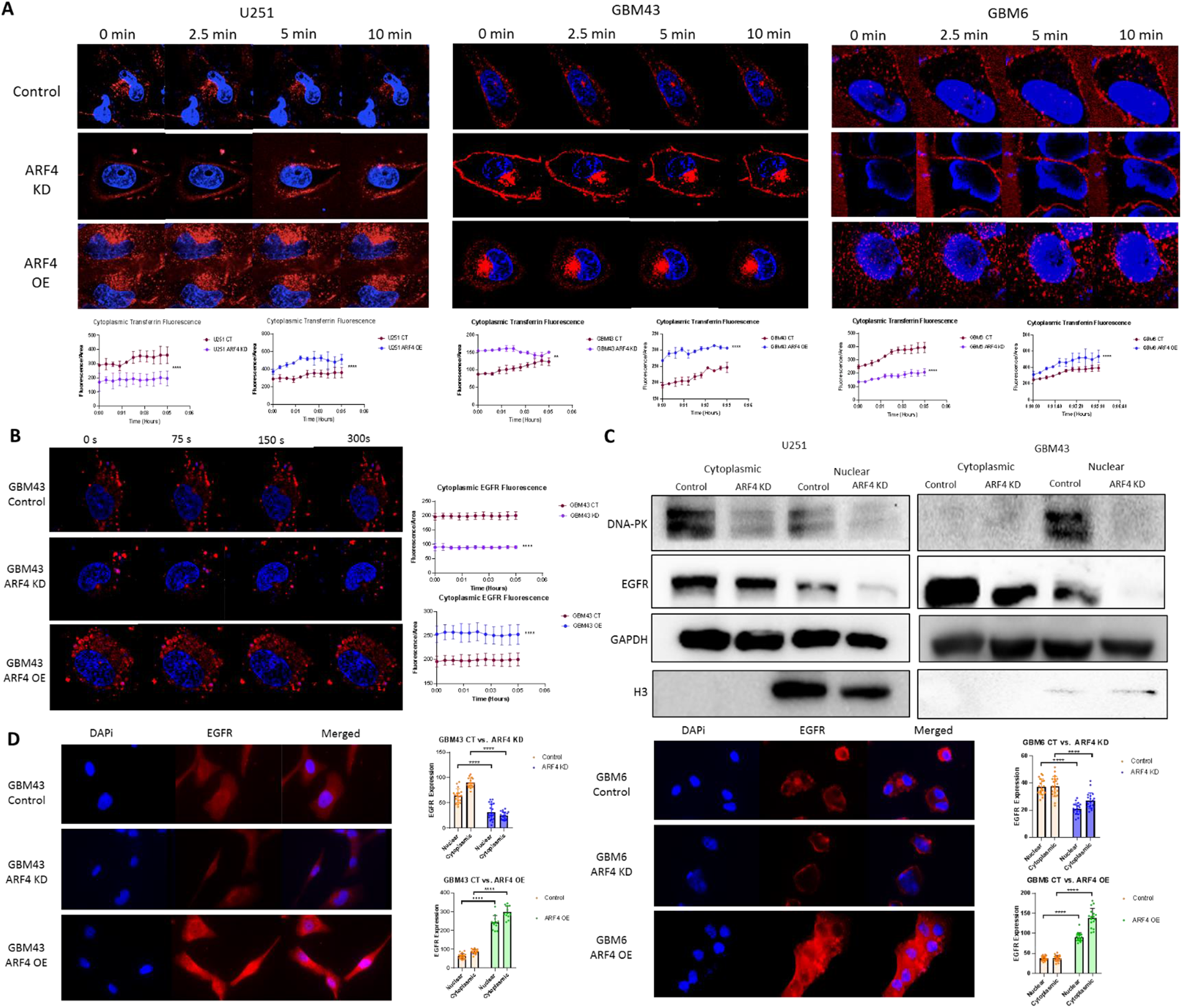
ARF4-mediated Retrograde Trafficking of EGFR to the Nucleus Promotes Chemoresistance through Transcriptional Activation of DNA-PK. ***A)*** Live cell imaging of transferrin receptor after 0, 2.5, 5, and 10 minutes of exogenous introduction in control, ARF4-knockdown, and ARF4-overexpression U251, GBM43, and GBM6 cells, along with quantitative analysis of cytoplasmic fluorescence over time. ***B)*** Live cell imaging of EGFR after 0, 75, 150, and 300 seconds of EGF introduction in control, ARF4-knockdown, and ARF4-overexpression GBM43 cells, along with quantitative analysis of cytoplasmic fluorescence over time. ***C)*** Immunoblot analysis of *EGFR* and *DNA-PK* in cytoplasmic and nuclear fractions of control, ARF4-knockdown, and ARF4-overexpression GBM43 cells. Protein extract of these lines were immunoblotted with antibody against *EGFR* and *DNA-PK* or an antibody against *GAPDH* as a control for equal loading of cytoplasmic protein and *H3* for nuclear protein. ***D)*** Immunocytochemistry of EGFR in control, ARF4-knockdown, and ARF4-overexpression GBM43 and GBM6 cells, along with quantitative analysis of nuclear vs. cytoplasmic fluorescence. Analysis was performed in Prism 8, using ANOVA to compare row-means to determine significance or using log-rank tests to determine survival significance *p < 0.05; **p < 0.01; ***p < 0.001; ****p < 0.0001; ns, not significant.

After imaging the effects of *ARF4*-knockdown and *ARF4*-overexpression on retrograde transport, *EGFR* trafficking was then recorded and measured, showing expected results of significantly less cytoplasmic fluorescence in the *ARF4*-knockdowns *(p<0.0001)* and significantly more cytoplasmic fluorescence in the *ARF4*-overexpressions *(p<0.0001)* when compared to control cells **(Fig 6B)**. We were thus able to verify that the changes in *EGFR* trafficking are due to an *ARF4*-dependent mechanism by which receptor mislocalization to the nucleus drives chemoresistance.

We then validated these trends by measuring expression of retrograde transport marker *EHD3* as well as RTKs *EGFR* and *VEGFR* via western blotting. Western blots comparing control to *ARF4*-knockdown conditions revealed that there was not only lower expression of *EHD3*, but also lower expression of RTKs previously investigated and *DNA-PK* across two cell lines **(Fig S5C)**. Further investigation after nuclear and cytoplasmic fractionation also demonstrated a drastic decrease in *EGFR* and *DNA-PK* expression in the nucleus when *ARF4* was knocked down **(Fig 6C)**. This was also seen from immunocytochemistry stainings comparing control GBM43 and GBM6 cells to *ARF4*-knockdown and *ARF4*-overexpression cells, revealing lower and higher nuclear *EGFR* expression, respectively *(p<0.0001)* **(Fig 6D)**.

Up until now, we have linked *EGFR* mislocalization during TMZ treatment to changes in *ARF4* upregulation. To fully confirm that this effect is due to *ARF4*’s role in retrograde trafficking specifically, we treated GBM43 cells with retrograde transport inhibitor Bafa1. We expected to see a trend similar to that of an *ARF4*-knockdown cell, where impaired trafficking results in lower *EGFR* transportation to the nucleus. Indeed, live cell imaging revealed reduced trafficking for *TfR*, where it stays packed on the cell membrane **(Fig S5D)**. Furthermore, western blot analysis reveals a similar decrease in nuclear *EGFR* expression **(Fig S5E)**.

We finally wanted to determine the impact of *ARF4*-knockdown and *ARF4*-overexpression on DNA damage marker γH2AX. *ARF4*-knockdown GBM43 and GBM6 cells resulted in dramatic increases in γH2AX foci *(p<0.0001)*, while ARF4-overexpression cells results in a slight decrease in γH2AX foci, substantiating the significance of *ARF4*’s downstream effects on *DNA-PK* transcriptional activation as a response to chemotherapy **(Fig S5F)**.

## Discussion

Glioblastoma is an aggressive and deadly cancer, with one of the worst survival rates among all human cancers. Despite ongoing advances in therapeutic modalities, GBM still remains incurable due to its reliable development of resistance to treatment. As such, this disease desperately requires novel, unbiased techniques to elucidate previously unknown drivers of this fatal resistance.

In this study, we perform a whole-genome CRISPR-Cas9 TMZ-sensitivity screen, tapping into its ability to expose genetic vulnerabilities from a functional standpoint. We identified approximately 6000 genes that were found to be critical for conferring resistance to TMZ; of these, 150 targets were deemed significant across two analytical methods. Although all of these genes represent promising targets for preventing GBM’s acquisition of resistance, we performed validation experiments on four previously unstudied genes, all of which reflected key pathways enriched in our screen. Patient data and in vitro assessment of expression and viability revealed that these genes do indeed play a role in driving GBM resistance to TMZ, further verifying the accuracy of our screen and cementing these genes as promising therapeutic targets.

One of these genes, *ARF4*, was of particular interest to us, due to its involvement in retrograde trafficking through regulation of endosomal formation. Although retrograde trafficking is normally a well-regulated process by which endosomes are transported to either the trans-Golgi network, the perinuclear region, or the nucleus, retrograde trafficking in cancer has been found to be highly dysregulated and even associated with enhanced proliferation and metastasis [27]. Despite its extensive study in cancer, there are still gaps in the literature around the mechanism by which retrograde trafficking may drive therapeutic resistance in any cancer, including GBM. Due to the lack of study in this area, we sought to determine the mechanism by which altered retrograde trafficking seen during therapy is able to drive GBM’s resistance.

Here, we show that TMZ treatment results in increased retrograde trafficking through ER stress-induced upregulation of *ARF4*. Further investigation revealed that one effect of this increased trafficking is the acceleration of RTK transport to the nucleus, which has already been linked extensively to chemoresistance [47]. More specifically, EGFR was seen to be retro-translocated into the nucleus at much greater levels. This higher expression of nuclear EGFR resulted in less DNA damage due to greater transcriptional activation of DNA-PK. Given nuclear EGFR‘s role in strengthening DNA repair response, this sheds important light on how GBM cells are able to avoid therapy-induced DNA damage over time and how retrograde trafficking plays such a key role in the receptor mislocalization that is evident in resistant tumors. Ultimately, we believe that this new understanding of retrograde trafficking has revealed ways in which GBM is able to develop resistance to therapy so quickly, revealing a unique means of preventing GBM’s deadly recurrence.

## Acknowledgments

This work was supported by the National Institute of Neurological Disorders and Stroke grant 1R01NS096376, 1R01NS112856 the American Cancer Society grant RSG-16-034-01-DDC (to A.U.A.) and P50CA221747 SPORE for Translational Approaches to Brain Cancer.

The results published here are in part based upon data generated by the TCGA Research Network: https://www.cancer.gov/tcga, and were further analyzed through GlioVis. In addition, these results use data generated by the Human Protein Atlas and GBMSeq (Gephart Lab, www.gbmseq.org). Figures, in part, were generated using BioRender (www.biorender.com).

## Author Contributions

Conceptualization: Shreya Budhiraja, Adam M Sonabend, Atique U Ahmed Methodology: Shreya Budhiraja, Shivani Baisiwala, Adam M Sonabend, Atique U Ahmed Validation, Formal Analysis, Investigation: Shreya Budhiraja, Shivani Baisiwala, Ella Perrault, Khizar Nandoliya, Sia Cho, Gabriel Dara, Andrew Zolp, Li Chen, Crismita Dmello, Cheol H Park

Resources: Atique U Ahmed, Adam M Sonabend

Data Curation & Draft Preparation: Shreya Budhiraja

Review & Editing: Shreya Budhiraja, Shivani Baisiwala, Atique U Ahmed, Adam M Sonabend

Supervision, Project Administration, Funding Acquisition: Atique U Ahmed, Adam M Sonabend

---

**Figure S1:**
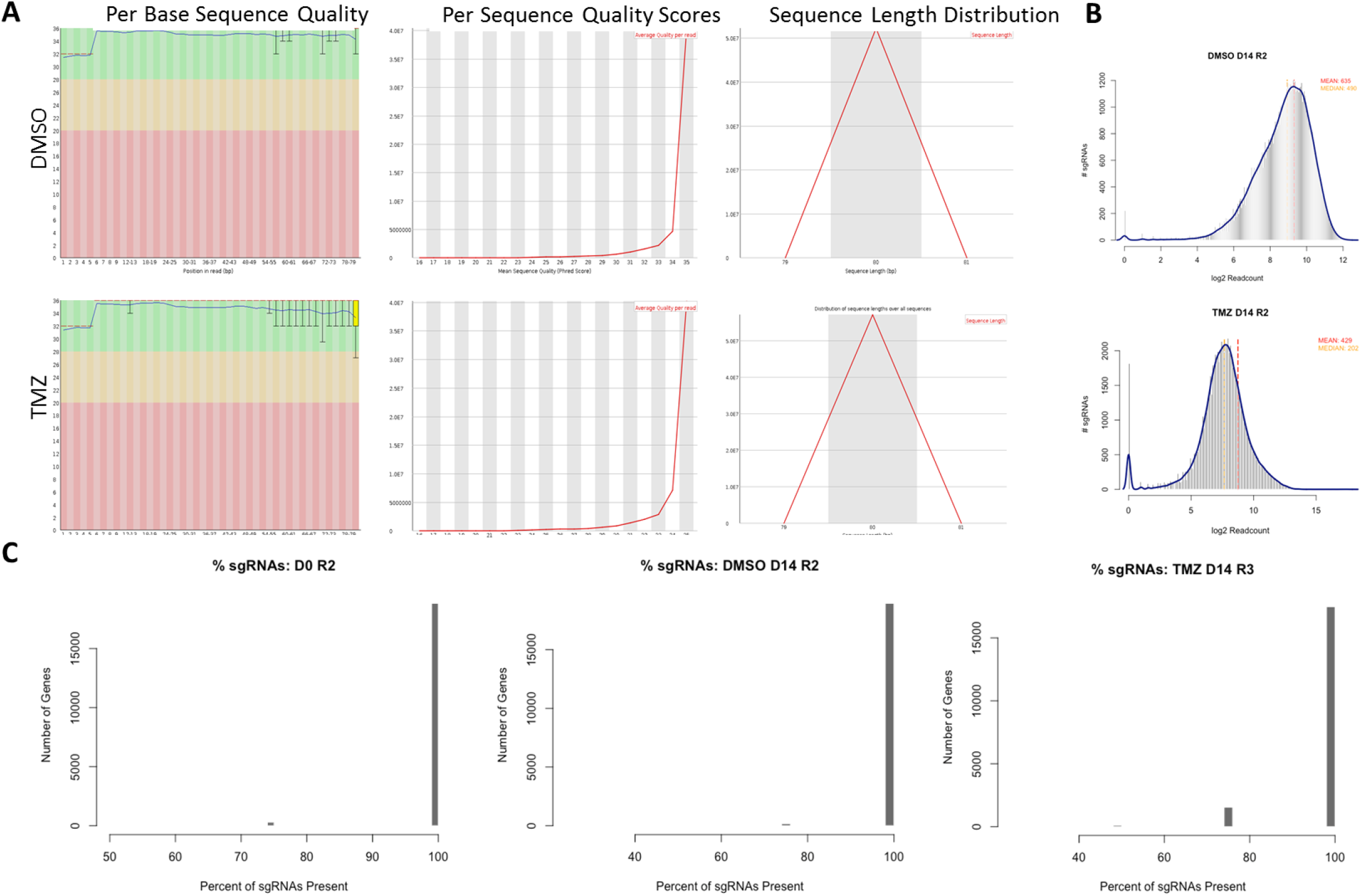
CRISPR-Cas9 Screen and Sequencing Quality. ***A)*** Per base sequence quality, per sequence quality scores, and sequence length distribution for DMSO and TMZ conditions at d14 are appropriate (B) Cumulative frequency at day 14 was assessed and reflected an appropriate distribution (C) Guide coverage reflected minimal guide loss at d14

**Figure S2:**
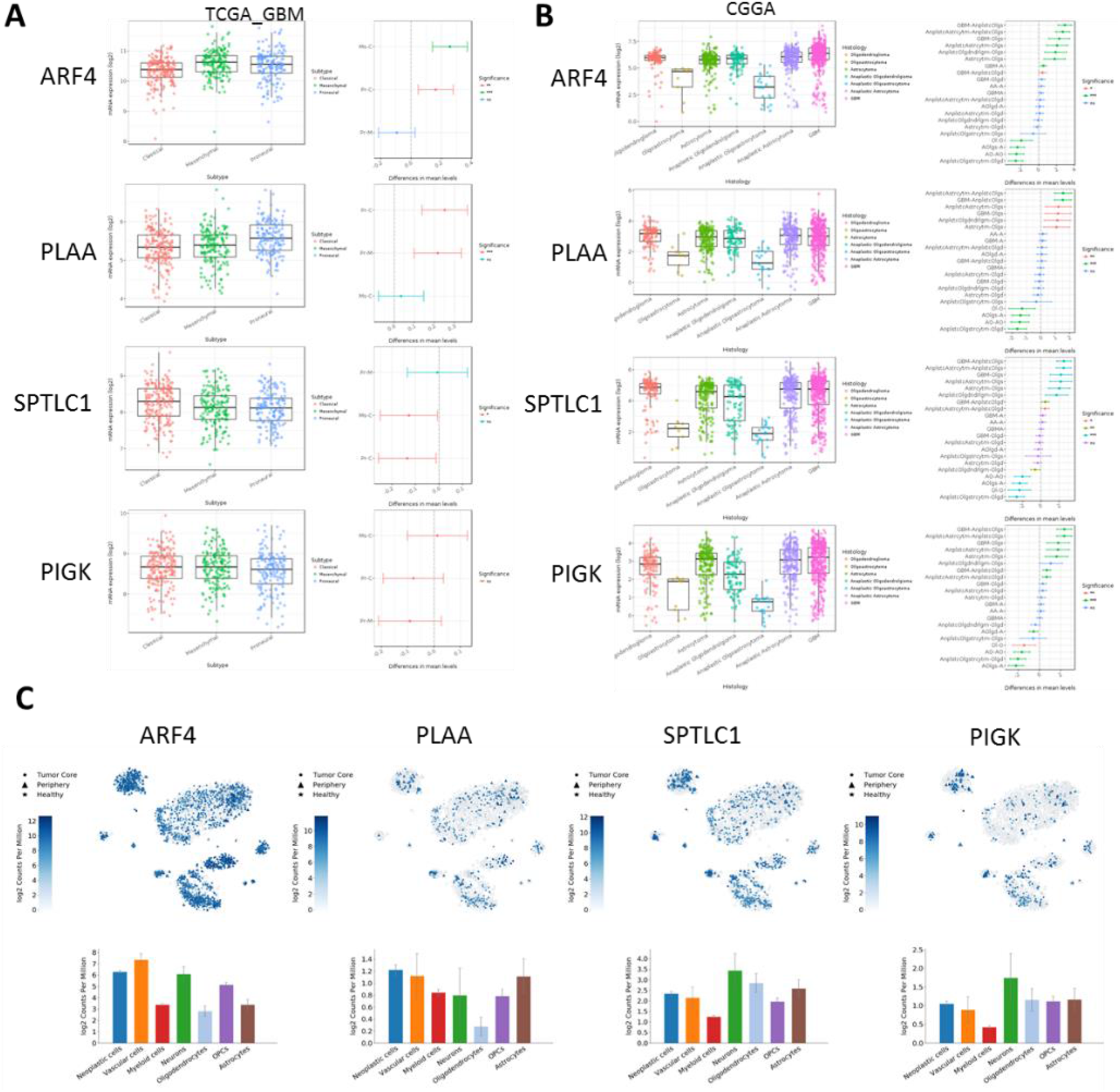
Target Gene Expression in Patient Datasets. ***A)** ARF4, PLAA, SPTLC1*, and *PIGK* mRNA expression levels were analyzed using the Cancer Genome Atlas (TCGA) for different GBM subtypes. ***B)*** *ARF4, PLAA, SPTLC1*, and *PIGK* mRNA expression levels were analyzed using the Cancer Genome Atlas (TCGA) for different glioma types based on aggressiveness. ***C)*** *ARF4, PLAA, SPTLC1*, and *PIGK* mRNA expression levels were analyzed using GBMSeq for expression among different cell types. Analysis was performed in Prism 8, using ANOVA to compare row-means to determine significance or using log-rank tests to determine survival significance *p < 0.05; **p < 0.01; ***p < 0.001; ****p < 0.0001; ns, not significant.

**Figure S3:**
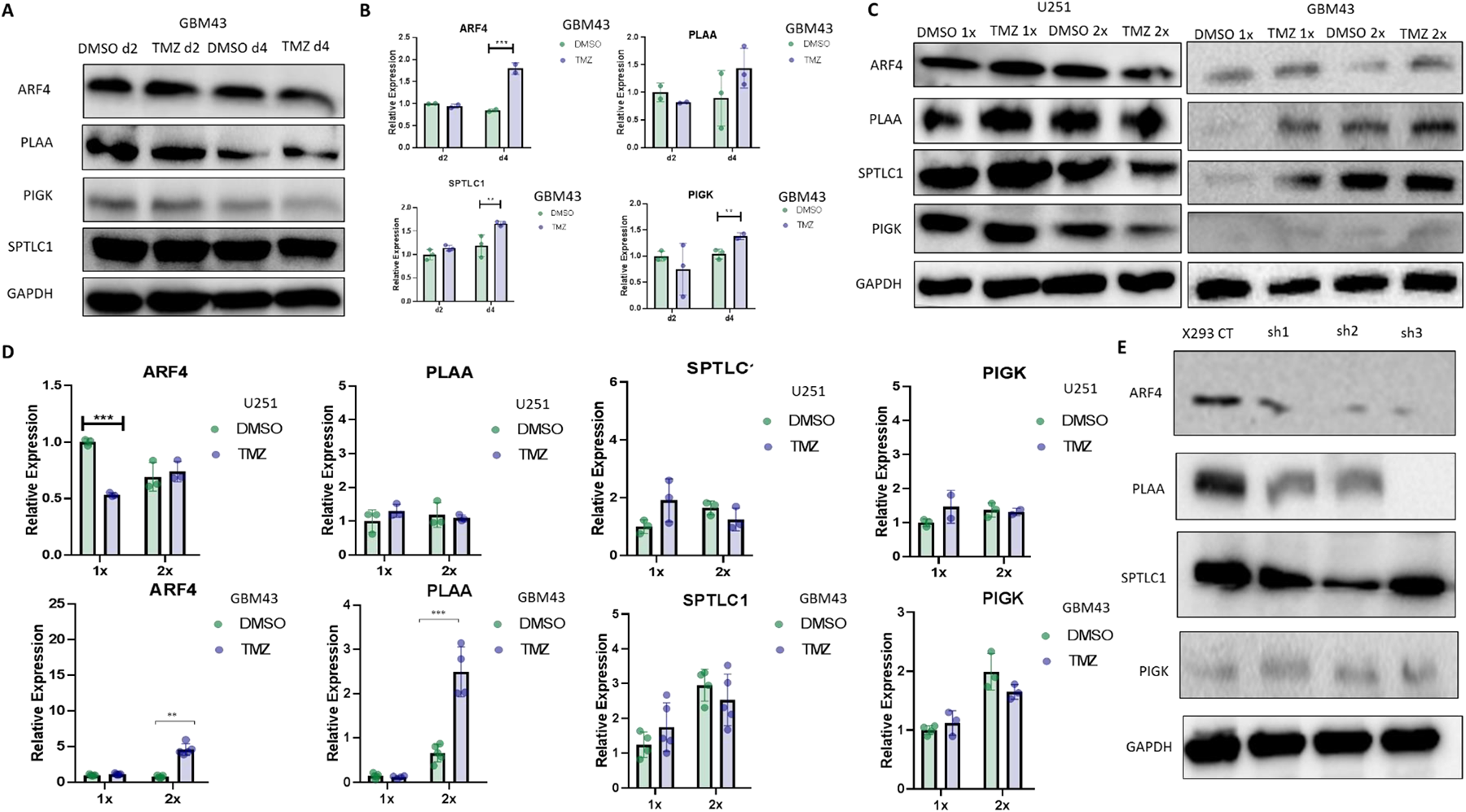
1x and 2x Treatment with TMZ does not Induce TMZ-resistance Gene Upregulation. ***A)*** Immunoblot analysis of *ARF4, PLAA, SPTLC1*, and *PIGK* protein expression in DMSO- and TMZ-treated GBM43 cells. Protein extract after 2 and 4 days of treatment were immunoblotted with antibody against *ARF4, PLAA, SPTLC1, and PIGK* or an antibody against *GAPDH* as a control for equal loading. ***B)*** *ARF4, PLAA, SPTLC1*, and *PIGK* mRNA expression levels were analyzed in DMSO- and TMZ-treated GBM43 cells using quantitative real-time PCR (qPCR). mRNA was harvested after 2 and 4 days of treatment. ***C)*** Immunoblot analysis of *ARF4, PLAA, SPTLC1*, and *PIGK* protein expression in 1x- and 2x-treated DMSO and TMZ U251 and GBM43 cells. Protein extract of these lines were immunoblotted with antibody against *ARF4, PLAA, SPTLC1, and PIGK* or an antibody against *GAPDH* as a control for equal loading. ***D)*** *ARF4, PLAA, SPTLC1*, and *PIGK* mRNA expression levels were analyzed in 1x- and 2x-DMSO and TMZ U251 and GBM43 cells using quantitative real-time PCR (qPCR). ***E)*** Immunoblot analysis of multiple shRNAs to knockdown targets of interest. Analysis was performed in Prism 8, using ANOVA to compare row-means to determine significance or using log-rank tests to determine survival significance *p < 0.05; **p < 0.01; ***p < 0.001; ****p < 0.0001; ns, not significant.

**Figure S4:**
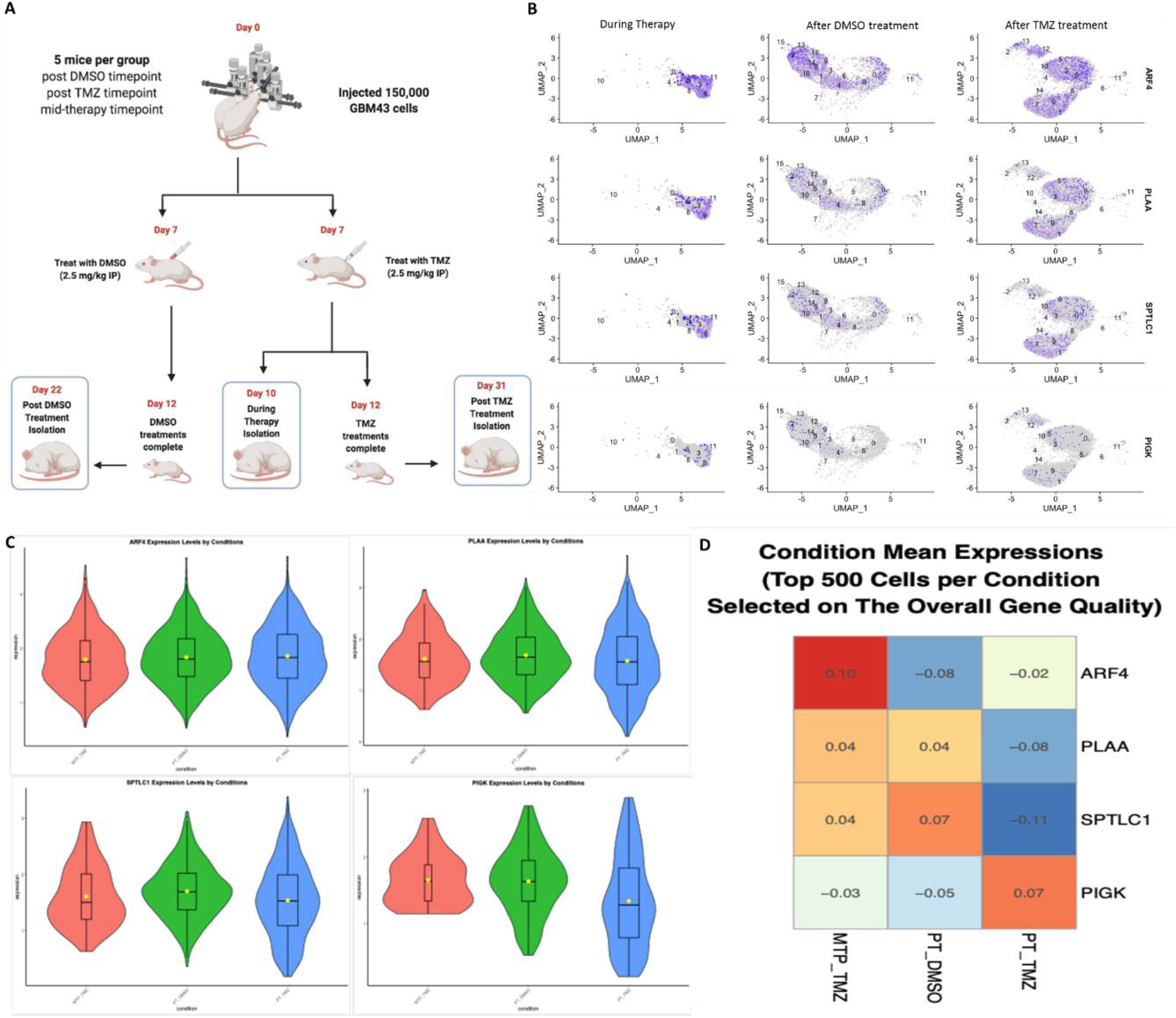
Single-cell RNA Sequencing Analysis for Genes of Interest Before, During, and After Therapy. ***A)*** Single-cell RNA Sequencing was performed. Three groups of mice were examined: one group was isolated post-DMSO treatment, one group was isolated post-TMZ treatment, and one group was isolated mid-TMZ treatment. Sequencing for *ARF4, PLAA, SPTLC1*, and *PIGK* RNA expression at these timepoints was performed. ***B)*** Feature plots of *ARF4, PLAA, SPTLC1*, and *PIGK* RNA expression distribution for each timepoint. ***C)*** Bar plots of *ARF4, PLAA, SPTLC1*, and *PIGK* mean RNA expression for each timepoint. ***D)*** Heat map of *ARF4, PLAA, SPTLC1*, and *PIGK* mean RNA expression.

**Figure S5:**
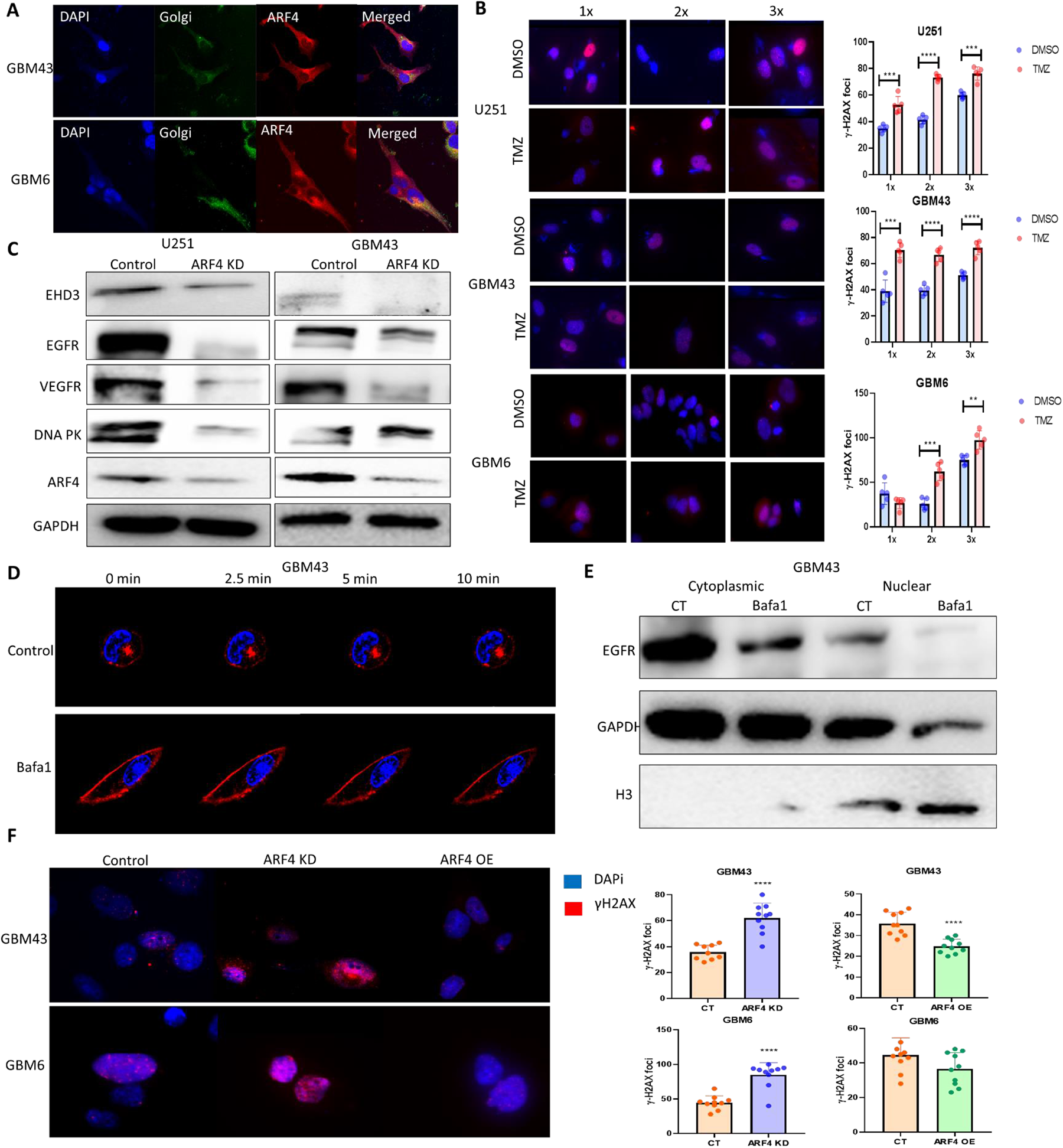
ARF4-mediated Retrograde Trafficking during TMZ Therapy. ***A)*** Immunocytochemistry of the Golgi apparatus and *ARF4* in GBM43 and GBM6 cells. ***B)*** Immunocytochemistry of the DNA damage marker γH2AX in 1x, 2x, and 3x DMSO- and TMZ-tread U251, GBM43, and GBM6 cells, along with quantitative analysis of γH2AX foci counts. ***C)*** Immunoblot analysis of *EHD3, EGFR, VEGFR*, and *DNA-PK* protein expression in control and ARF4-knockdown U251 and GBM43 cells. Protein extract of these lines were immunoblotted with antibody against *EHD3, EGFR, VEGFR*, and *DNA-PK* or an antibody against *GAPDH* as a control for equal loading. ***D)*** Live cell imaging of transferrin receptor after 0, 2.5, 5, and 10 minutes of exogenous introduction in control and Bafa1-(retrograde transport inhibitor) treated GBM43 cells. ***E)*** Immunoblot analysis of *EGFR* in cytoplasmic and nuclear fractions of control and Bafa1-treated GBM43 cells. Protein extract of these lines were immunoblotted with antibody against *EGFR* or an antibody against *GAPDH* as a control for equal loading of cytoplasmic protein and *H3* for nuclear protein. ***F)*** Immunocytochemistry of the DNA damage marker γH2AX in control, *ARF4*-knockdown, and *ARF4*-overexpression GBM43 and GBM6 cells, along with quantitative analysis of γH2AX foci counts. Analysis was performed in Prism 8, using ANOVA to compare row-means to determine significance or using log-rank tests to determine survival significance *p < 0.05; **p < 0.01; ***p < 0.001; ****p < 0.0001; ns, not significant.

## Notes

### Competing Interest Statement

The authors have declared no competing interest.

## References

1. S.P. Caragher, et al., of Dopamine Receptor 2 Prompts Transcriptomic and Metabolic Plasticity in Glioblastoma. The Journal of neuroscience: the official journal of the Society for Neuroscience, 2019. 39(11): p. 1982–1993.

2. Stupp, R., et al., Radiotherapy plus concomitant and adjuvant temozolomide for glioblastoma. N Engl J Med, 2005. 352(10): p. 987–96.

3. Lukas, R.V., et al., Newly Diagnosed Glioblastoma: A Review on Clinical Management. Oncology (Williston Park), 2019. 33(3): p. 91–100.

4. Baisiwala, S., et al., Chemotherapeutic Stress Induces Transdifferentiation of Glioblastoma Cells to Endothelial Cells and Promotes Vascular Mimicry. Stem cells international, 2019. 2019: p. 6107456–6107456.

5. Baumert, B.G., et al., Temozolomide chemotherapy versus radiotherapy in high-risk low-grade glioma (EORTC 22033-26033): a randomised, open-label, phase 3 intergroup study. The Lancet. Oncology, 2016. 17(11): p. 1521–1532.

6. Hottinger, A.F., P. Pacheco, and R. Stupp, Tumor treating fields: a novel treatment modality and its use in brain tumors. Neuro-oncology, 2016. 18(10): p. 1338–1349.

7. Doudna, J.A. and E. Charpentier, The new frontier of genome engineering with CRISPR-Cas9. Science, 2014. 346(6213): p. 1258096.

8. Joung, J., et al., Genome-scale CRISPR-Cas9 knockout and transcriptional activation screening. Nature protocols, 2017. 12(4): p. 828–863.

9. Adli, M., The CRISPR tool kit for genome editing and beyond. Nature Communications, 2018. 9(1): p. 1911.

10. Klinger SC, Siupka P, Nielsen MS. Retromer-mediated trafficking of transmembrane receptors and transporters. Membranes. 2015 Sep; 5(3):288–306.

11. Johannes L, Popoff V. Tracing the retrograde route in protein trafficking. Cell. 2008 Dec 26; 135(7):1175–87.

12. Alberts B, Johnson A, Lewis J, Raff M, Roberts K, Walter P. Transport from the trans Golgi network to lysosomes. InMolecular Biology of the Cell. 4th edition 2002. Garland Science.

13. Tokarev AA, Alfonso A, Segev N. Overview of Intracellular Compartments and Trafficking Pathways. InMadame Curie Bioscience Database [Internet] 2013. Landes Bioscience.

14. Awah, C.U., et al., Ribosomal protein S11 influences glioma response to TOP2 poisons. Oncogene, 2020. 39(27): p. 5068–5081.

15. Li, W., et al., MAGeCK enables robust identification of essential genes from genome-scale CRISPR/Cas9 knockout screens. Genome Biol, 2014. 15(12): p. 554.

16. Bageritz, J. and G. Raddi, Single-Cell RNA Sequencing with Drop-Seq, in Single Cell Methods: Sequencing and Proteomics, V. Proserpio, Editor. 2019, Springer New York: New York, NY. p. 73–85.

17. Stuart, T., et al., Comprehensive Integration of Single-Cell Data. Cell, 2019. 177(7): p. 1888–1902.e21.

18. Darmanis, S., et al., Single-Cell RNA-Seq Analysis of Infiltrating Neoplastic Cells at the Migrating Front of Human Glioblastoma. Cell reports, 2017. 21(5): p. 1399–1410.

19. Shinsato Y, Furukawa T, Yunoue S, Yonezawa H, Minami K, Nishizawa Y, Ikeda R, Kawahara K, Yamamoto M, Hirano H, Tokimura H. Reduction of MLH1 and PMS2 confers temozolomide resistance and is associated with recurrence of glioblastoma. Oncotarget. 2013 Dec; 4(12):2261.

20. Yip S, Miao J, Cahill DP, Iafrate AJ, Aldape K, Nutt CL, Louis DN. MSH6 mutations arise in glioblastomas during temozolomide therapy and mediate temozolomide resistance. Clinical cancer research. 2009 Jul 15; 15(14):4622–9.

21. McFaline-Figueroa JL, Braun CJ, Stanciu M, Nagel ZD, Mazzucato P, Sangaraju D, Cerniauskas E, Barford K, Vargas A, Chen Y, Tretyakova N. Minor changes in expression of the mismatch repair protein MSH2 exert a major impact on glioblastoma response to temozolomide. Cancer research. 2015 Aug 1; 75(15):3127–38.

22. Memon AR. The role of ADP-ribosylation factor and SAR1 in vesicular trafficking in plants. Biochimica et Biophysica Acta (BBA)-Biomembranes. 2004 Jul 1; 1664(1):9–30.

23. Jackson CL. Arf proteins and their regulators: at the interface between membrane lipids and the protein trafficking machinery. InRas superfamily small G proteins: biology and mechanisms 2 2014 (pp. 151–180). Springer, Cham.

24. Nakai W, Kondo Y, Saitoh A, Naito T, Nakayama K, Shin HW. ARF1 and ARF4 regulate recycling endosomal morphology and retrograde transport from endosomes to the Golgi apparatus. Molecular biology of the cell. 2013 Aug 15; 24(16):2570–81.

25. Ma M, Burd CG. Retrograde trafficking and plasma membrane recycling pathways of the budding yeast Saccharomyces cerevisiae. Traffic. 2020 Jan; 21(1):45–59.

26. Spang A. Retrograde traffic from the Golgi to the endoplasmic reticulum. Cold Spring Harbor perspectives in biology. 2013 Jun 1; 5(6):a013391.

27. Maisel SA, Schroeder J. Wrong place at the wrong time: how retrograde trafficking drives cancer metastasis through receptor mislocalization. Journal of Cancer Metastasis and Treatment. 2019 Feb 13;5.

28. Howley BV, Howe PH. Metastasis-associated upregulation of ER-Golgi trafficking kinetics: regulation of cancer progression via the Golgi apparatus. Oncoscience. 2018 May; 5(5-6):142.

29. Tu Y, Zhao L, Billadeau DD, Jia D. Endosome-to-TGN trafficking: organelle-vesicle and organelle-organelle interactions. Frontiers in cell and developmental biology. 2020 Mar 18; 8:163.

30. Chen MK, Du Y, Sun L, Hsu JL, Wang YH, Gao Y, Huang J, Hung MC. H2O2 induces nuclear transport of the receptor tyrosine kinase c-MET in breast cancer cells via a membrane-bound retrograde trafficking mechanism. Journal of Biological Chemistry. 2019 May 24; 294(21):8516–28.

31. Reiling JH, Olive AJ, Sanyal S, Carette JE, Brummelkamp TR, Ploegh HL, Starnbach MN, Sabatini DM. A CREB3–ARF4 signalling pathway mediates the response to Golgi stress and susceptibility to pathogens. Nature cell biology. 2013 Dec; 15(12):1473–85.

32. Van Raam BJ, Lacina T, Lindemann RK, Reiling JH. Secretory stressors induce intracellular death receptor accumulation to control apoptosis. Cell death & disease. 2017 Oct; 8(10):e3069-.

33. Sampieri L, Di Giusto P, Alvarez C. CREB3 transcription factors: ER-golgi stress transducers as hubs for cellular homeostasis. Frontiers in cell and developmental biology. 2019 Jul 3; 7:123.

34. Liu Z, Lv Y, Zhao N, Guan G, Wang J. Protein kinase R-like ER kinase and its role in endoplasmic reticulum stress-decided cell fate. Cell death & disease. 2015 Jul; 6(7):e1822.

35. Oslowski CM, Urano F. Measuring ER stress and the unfolded protein response using mammalian tissue culture system. Methods in enzymology. 2011 Jan 1; 490:71–92.

36. Guérin R, Arseneault G, Dumont S, Rokeach LA. Calnexin is involved in apoptosis induced by endoplasmic reticulum stress in the fission yeast. Molecular biology of the cell. 2008 Oct; 19(10):4404–20.

37. Nirschl JJ, Holzbaur EL. Live-cell imaging of retrograde transport initiation in primary neurons. Methods in cell biology. 2016 Jan 1; 131:269–76.

38. York HM, Patil A, Moorthi UK, Kaur A, Bhowmik A, Hyde GJ, Gandhi H, Fulcher A, Gaus K, Arumugam S. Rapid whole cell imaging reveals a calcium-APPL1-dynein nexus that regulates cohort trafficking of stimulated EGF receptors. Communications biology. 2021 Feb 17; 4(1):1–3.

39. Van Dam EM, Stoorvogel W. Dynamin-dependent transferrin receptor recycling by endosome-derived clathrin-coated vesicles. Molecular biology of the cell. 2002 Jan 1; 13(1):169–82.

40. Mayle KM, Le AM, Kamei DT. The intracellular trafficking pathway of transferrin. Biochimica et Biophysica Acta (BBA)-General Subjects. 2012 Mar 1; 1820(3):264–81.

41. Taeger J, Moser C, Hellerbrand C, Mycielska ME, Glockzin G, Schlitt HJ, Geissler EK, Stoeltzing O, Lang SA. Targeting FGFR/PDGFR/VEGFR impairs tumor growth, angiogenesis, and metastasis by effects on tumor cells, endothelial cells, and pericytes in pancreatic cancer. Molecular cancer therapeutics. 2011 Nov 1; 10(11):2157–67.

42. Liccardi G, Hartley JA, Hochhauser D. EGFR nuclear translocation modulates DNA repair following cisplatin and ionizing radiation treatment. Cancer research. 2011 Feb 1; 71(3):1103–14.

43. Friedmann BJ, Caplin M, Savic B, Shah T, Lord CJ, Ashworth A, Hartley JA, Hochhauser D. Interaction of the epidermal growth factor receptor and the DNA-dependent protein kinase pathway following gefitinib treatment. Molecular cancer therapeutics. 2006 Feb 1; 5(2):209–18.

44. Annovazzi LA, Caldera V, Mellai M, Riganti C, Battaglia L, Chirio D, Melcarne A, Schiffer D. The DNA damage/repair cascade in glioblastoma cell lines after chemotherapeutic agent treatment. International journal of oncology. 2015 Jun 1; 46(6):2299–308.

45. Tomas A, Futter CE, Eden ER. EGF receptor trafficking: consequences for signaling and cancer. Trends in cell biology. 2014 Jan 1; 24(1):26–34.

46. Fraser J, Simpson J, Fontana R, Kishi-Itakura C, Ktistakis NT, Gammoh N. Targeting of early endosomes by autophagy facilitates EGFR recycling and signalling. EMBO reports. 2019 Oct 4; 20(10):e47734.

47. Carrasco-García E, Saceda M, Martínez-Lacaci I. Role of receptor tyrosine kinases and their ligands in glioblastoma. Cells. 2014 Jun; 3(2):199–235.

